# Systematic Phenotyping and Characterization of the 5xFAD mouse model of Alzheimer’s Disease

**DOI:** 10.1101/2021.02.17.431716

**Authors:** Stefania Forner, Shimako Kawauchi, Gabriela Balderrama-Gutierrez, Enikö A. Kramár, Dina P. Matheos, Jimmy Phan, Dominic I. Javonillo, Kristine Minh Tran, Edna Hingco, Celia da Cunha, Narges Rezaie, Joshua Andrei Alcantara, David Baglietto-Vargas, Camden Jansen, Jonathan Neumann, Marcelo A. Wood, Grant R. MacGregor, Ali Mortazavi, Andrea J. Tenner, Frank M. LaFerla, Kim N. Green

## Abstract

Mouse models of human diseases are invaluable tools for studying pathogenic mechanisms and testing interventions and therapeutics. For disorders such as Alzheimer’s disease in which numerous models are being generated, a challenging first step is to identify the most appropriate model and age to effectively evaluate new therapeutic approaches. Here we conducted a detailed phenotypic characterization of the 5xFAD model on a congenic C57BL/6J strain background, across its lifespan – including a seldomly analyzed 18-month old time point to provide temporally correlated phenotyping of this model and a template for characterization of new models of LOAD as they are generated. This comprehensive analysis included quantification of plaque burden, Aβ biochemical levels, and neuropathology, neurophysiological measurements and behavioral and cognitive assessments, and evaluation of microglia, astrocytes, and neurons. Analysis of transcriptional changes was conducted using bulk-tissue generated RNA-seq data from microdissected cortices and hippocampi as a function of aging, which can be explored at the UCI Mouse Explorer and AD Knowledge Portal. This deep-phenotyping pipeline identified novel aspects of age-related pathology in the 5xFAD model.

## Background and Summary

Animal models of Alzheimer’s disease play a pivotal role in facilitating our understanding of disease mechanism and for drug discovery. Yet, despite their promise, there has been significant concern about their translational reliability, particularly as treatments effective in mouse models have largely proven ineffectual when evaluated in clinical trials^1-3^. Several factors likely underlie these translational failures, but two prominent reasons are that the vast majority of AD animal models are based on overexpression and the inclusion of autosomal dominant mutations, despite the fact overexpression or genetic mutations do not occur in the overwhelming majority of human AD cases.

In 2015 the US NIH/NIA initiated a new program called Model Organism Development and Evaluation for Late-Onset Alzheimer’s Disease (MODEL-AD; https://www.model-ad.org/) to develop the next generation of animal models. MODEL-AD specifically seeks to better recapitulate the etiology and mechanisms of late-onset Alzheimer’s disease (LOAD), with the ultimate goal of improving translatability. Accomplishing this ambitious objective requires a multiprong strategy, including, in addition to the generation of models more aligned with LOAD, the detailed development of a standardized characterization of a phenotyping pipeline that can provide comprehensive comparative data about molecular, cellular and functional changes that occur as a function of age and brain region. As part of this goal, it is critical that established AD mouse models serve as a benchmark for future comparisons. The 5xFAD mouse (Tg(APPSwFlLon,PSEN1*M146L*L286V)6799Vas/Mmjax) was generated in 2006^4^ and displays overexpression of APP and PSEN1 containing 5 familial AD mutations (APP KM670/671NL (Swedish), APP I716V (Florida), APP V717I (London), PSEN1 M146L, PSEN1 L286V), under the control of a *Thy1* mini-gene^5,6^, which directs expression to forebrain neurons. 5xFAD mice develop robust amyloid pathologies, with plaques appearing in the brain from 2-4 months of age^7^, triggering robust microgliosis and inflammatory processes^4,7^ as well as synaptic^7^ and neuronal loss^4,7^. Because the 5xFAD mouse is commonly used— ∼10% of all AD studies that use an animal model employ this strain (AlzPED; https://alzped.nia.nih.gov/), we included it as a benchmark reference model for our studies. Here, we used the 5xFAD mouse model to evaluate our deep-phenotyping pipeline, including an 18-month time point which is rarely analyzed by researchers in the AD field. Through the evaluation of behavior and cognition, long-term potentiation, gene expression, among other parameters across the lifespan (4, 8, 12, and 18 months of age), we demonstrate that the analytical pipeline used provides robust information relevant to understand changes that occur during development of pathology in a mouse model of AD. This data is freely accessible to the public through the AD Knowledge Portal (https://adknowledgeportal.synapse.org/) and should prove useful to AD investigators.

## Methods

### Animals

All animal experiments were approved by the UC Irvine Institutional Animal Care and Use Committee and were conducted in compliance with all relevant ethical regulations for animal testing and research. 5xFAD hemizygous (B6.Cg-Tg(APPSwFlLon,PSEN1*M146L*L286V)6799Vas/Mmjax, Stock number 34848-JAX, MMRRC) and its wildtype littermates were produced by crossing or IVF procedures with C57BL/6J (Jackson Laboratory, ME) females. After weaning, they were housed together with littermates and aged until the harvest dates. For 5xFAD genotyping, Hydrolysis probe which hybridizes APP(swe) mutation amplicon was used (For 5’-TGGGTTCAAACAAAGGTGCAA -3’ and Rev 5’-GATGACGATCACTGTCGCTATGAC-3’: APP(swe) probe 5’-CATTGGACTCATGGTGGGCGGTG-3’.) to detect transgenes. We used the endogenous Apo B allele (For 5’-CACGTGGGCTCCAGCATT-3’ and Rev 5’-TCACCAGTCATTTCTGCCTTTG-3’: ApoB probe 5’-CCAATGGTCGGGCACTGCTCAA-3’) to normalize the Ct values. All animals were bred by the Transgenic Mouse Facility at UCI.

### Behavioral Testing

Noldus Ethovision software (Wageningen, Netherlands) was employed to video-record and track animal behavior and analyses were performed by Ethovision software. All protocols are publicly available through the AD Knowledge Portal (https://adknowledgeportal.synapse.org/) and the following behavioral paradigms were carried out according to established protocol^4,8,9^ and described briefly below:

#### Elevated plus maze (EPM)

Mice were placed in the center of an elevated plus maze (arms 6.2 x 75cm, with side walls 20 cm high on two closed arms, elevated 63 cm above the ground) for 5 min to assess anxiety. Automated scoring assessed the amount of time each mouse spent cumulatively in the open arms and closed arms of the maze.

#### Open field (OF)

In brief, mice were placed in a white box (33.7 cm L x 27.3 cm W x 21.6 cm H) for 5 min to assess motor function and anxiety and videotaped for 5 min. Videos were scored for % time in center of arena, distance traveled and speed.

#### Contextual Fear conditioning (CFC)

Behavior was scored using Noldus Ethovision v.14.0.1322. Activity Analysis to detect activity levels and freezing behaviors for both training and testing sessions. Each of the four CFC chambers (Ugo Basile, Germany) is inside a sound-attenuating boxes with ventilating fan, a dual (visible/I.R.) light, a speaker and a USB-camera. Each FC-Unit has an individual controller on-board. The CFC chamber is cleaned at the start of testing and between every mouse with Ethanol 70% and paper towels to eliminate olfactory cues. In the training trial, each mouse is placed in the chamber for 2 min to allow for habituation and exploration of the context, after which a shock is applied for 3 s at 0.5 mA. The mice are returned to their cages after 30 s. Twenty-four hours later, testing was conducted, whereby animals were placed in the chamber to explore for 5 min. Sessions are recorded and immobility time is determined using EthoVision software.

#### Rotarod

Motor performance and motor learning were tested using the rotarod (Ugo Basile, Germany). Each mouse is weighed prior to testing. There are 6 lanes on the Rotarod, therefore 6 mice can be tested at once. Each group of 6 mice will be tested 5 times, for 5 min maximum (300 s) for each trial. Latency to fall served as an indicator of motor coordination.

### Hippocampal slice preparation and LTP recording

Hippocampal slices were prepared from 5xFAD (5 females and 5 males) and WT (5 females and 5 males) at 4, 8 and 12 months of age. Following isoflurane anesthesia, mice were decapitated and the brain was quickly removed and submerged in ice-cold, oxygenated dissection medium containing (in mM): 124 NaCl, 3 KCl, 1.25 KH_2_PO_4_, 5 MgSO_4_, 0 CaCl_2_, 26 NaHCO_3_, and 10 glucose. Coronal hippocampal slices (320 µm) were prepared using a Leica vibrating tissue slicer (Model: VT1000S) before being transferred to an interface recording containing preheated artificial cerebrospinal fluid (aCSF) of the following composition (in mM): 124 NaCl, 3 KCl, 1.25 KH_2_PO_4_, 1.5 MgSO_4_, 2.5 CaCl_2_, 26 NaHCO_3_, and 10 glucose and maintained at 31 ± 1°C. Slices were continuously perfused with this solution at a rate of 1.75-2 ml/min while the surface of the slices were exposed to warm, humidified 95% O_2_ / 5% CO_2_. Recordings began following at least 2 hours of incubation.

Field excitatory postsynaptic potentials (fEPSPs) were recorded from CA1b stratum radiatum using a single glass pipette filled with 2M NaCl (2-3 MΩ) in response to orthodromic stimulation (twisted nichrome wire, 65 µm diameter) of Schaffer collateral-commissural projections in CA1 stratum radiatum. In some slices two stimulation electrodes were used (positioned at sites CA1a and CA1c) to stimulate independent populations of synapses (experimental and control pathways) on CA1b pyramidal cells. Pulses were administered in an alternating fashion to the two electrodes at 0.03 Hz using a current that elicited a 50% maximal response. Paired-pulse facilitation was measured at 40, 100, and 200 sec intervals prior to setting baseline. After establishing a 10-20 minutes stable baseline, the orthodromic stimulated pathway was used to induce long-term potentiation (LTP) by delivering 5 ‘theta’ bursts, with each burst consisting of four pulses at 100 Hz and the bursts themselves separated by 200 msec (i.e., theta burst stimulation or TBS). The stimulation intensity was not increased during TBS. The control pathway was used to monitor for baseline drifts in the slice. Data were collected and digitized by NAC 2.0 Neurodata Acquisition System (Theta Burst Corp., Irvine, CA) and stored on a disk.

### Histology

Mice were euthanized at 4, 8, 12 and 18 months via CO_2_ inhalation and transcardially perfused with 1X phosphate buffered saline (PBS). For all studies, brains were removed, and hemispheres separated along the midline. Brain halves were either flash frozen for subsequent biochemical analysis or drop-fixed in 4% paraformaldehyde (PFA (Thermo Fisher Scientific, Waltham, MA)) for immunohistochemical analysis. Fixed half brains were sliced at 40 μm using a Leica SM2000R freezing microtome. □ All brain hemispheres have been processed and every 12^th^ brain slice imaged via a Zeiss Slidescanner using a 10X objective. Images were corrected for shading, stitched together, and exported for quantification in Bitplane Imaris Software. The following analyses were then performed.

### Immunofluorescence

For Thioflavin-S (Thio-S) staining, free-floating sections were washed with 1X PBS (1×5 min), followed by dehydration in a graded series of ethanol (100%, 95%, 70%, 50%; 1×3 min each). The sections were incubated in 0.5% Thio-S (in 50% ethanol, Sigma-Aldrich) for 10 min. This was followed by 3×5 min washes with 50% ethyl alcohol and a final wash in 1X PBS (1×10 min). For 6E10 immunohistochemistry, sections were briefly rinsed in 1X PBS (1×5 min) followed by 10 min wash in 1X PBS. Following Thio-S staining or formic acid pretreatment (if required), sections underwent a standard indirect immunohistochemical protocol. To that end, free-floating sections were washed with 1X PBS (1×5 min), and immersed in normal serum blocking solution (5% normal goat serum with 0.2% Triton-X100 in 1X PBS) for 60 min. Primary antibodies and dilutions used are as follows: anti-ionized calcium-binding adapter molecule 1 (IBA1; 1:2000; 019-19741; Wako, Osaka, Japan), anti-Aβ_1-16_ (6E10; 1:2000; 803001; BioLegend, San Diego, CA), anti-S100β (1:200; ab41548; Abcam, Cambridge, MA), anti-glial fibrillary protein (GFAP; 1:1000; ab134436; Abcam), anti-Fox 3 protein (NeuN; 1:1000; ab104225; Abcam), anti-Ctip2 (CTIP2; 1:300; ab18465; Abcam), anti-lysosome::Jassociated membrane protein 1 (LAMP1; 1:200; ab25245; Abcam) and Thioflavin-S (0.5% ThioS in 50% Ethanol; Sigma-Aldrich).

### Imaris Quantitative Analysis

Volumetric image measurements were made in the hippocampus using Imaris software (Bitplane Inc.). Amyloid burden was acquired by measuring the total number of Aβ plaques and their size, expressed in area units (µm2) in the whole hippocampal area analyzed in an individual section. The 6E10-immunopositive signal (Aβ plaques) within the selected brain region was identified by a threshold level mask, which was maintained throughout the whole analysis per timeframe for uniformity. The total number of amyloid plaques and their area was obtained automatically by Imaris software. Quantitative comparisons between groups were always carried out on comparable sections of each animal processed at the same time with same batches of solutions. Microglial and astroglial loads (Iba1/GFAP-immunopositive) were counted with Bitplane Imaris software and normalized to the area of the hippocampus, subicullum, and cortex.

### Aβ soluble and insoluble fraction levels

The flash-frozen hemispheres of minimum 6 females and 6 males per age and per genotype were microdissected into cortical and hippocampal regions and then ground with a mortar and pestle to yield a homogenized tissue. One-half of the powder from the cortical region was homogenized in 1000 μl Tissue Protein Extraction Reagent (TPER) per 150mg and 150μl TPER for hippocampal region (Life Technologies, Grand Island, NY), respectively, with protease (Roche, Indianapolis, IN) and phosphatase inhibitors (Sigma-Aldrich, St. Louis, MO) and centrifuged at 100,000 g for 1 hour at 4°C to generate TPER-soluble fractions. For formic acid-fractions, pellets from TPER-soluble fractions were homogenized in 70% Formic Acid, half of TPER amount for cortical region and 75 μl for hippocampal region. Afterwards, the samples were centrifuged at 100,000 g for 1 hour at 4°C. Protein concentration in each fraction was determined via Bradford10,11.

Electrochemiluminescence-linked immunoassay Quantitative biochemical analyses of human Aβ soluble and insoluble fraction levels were performed using the V-PLEX Aβ Peptide Panel 1 (6E10) and (Meso Scale Discovery (MSD, Rockville MD, USA) according to the manufacturer’s instructions

### RNA Sequencing

Libraries were constructed by using the Nextera DNA Sample Preparation Kit (Illumina). Libraries were base-pair selected based on Agilent 2100 Bioanalyzer profiles and normalized determined by KAPA Library Quantification Kit (Illumina). The libraries were built from 5 different mice per genotype, sex and tissue (hippocampus and cortex) across 4 different timepoints (4, 8, 12 and 18 months). Sequences were aligned to the mouse genome (mm10) and annotation was done using GENCODE v21. Reads were mapped with STAR (2.5.1b-static) and RSEM (1.2.22) was used for quantification of gene expression.

#### Differential gene expression analysis

Differential gene expression analysis was done using edgeR12 per timepoint and tissue. Genes with an FDR > 0.05 were labeled. To compare different sets of genes differentially expressed we created a binary matrix identifying up and downregulated genes across different comparisons. A matrix indicating up or downregulation was later used to plot a heatmap.

From the comparisons, lists of genes of interest were chosen to plot a heatmap of their expression and a GO term enrichment analysis using enrichR (https://amp.pharm.mssm.edu/Enrichr/) and the top 5 GO terms were plotted. For comparing AMP-AD modules to 5xFAD gene lists obtained by edgeR, we calculated the fraction by counting the number of common genes between two gene lists and dividing by the number of genes in 5xFAD gene list for each comparison. We used Fisher exact test, as a procedure for obtaining exact probabilities associated with statistical hypotheses about 2X 2 contingency tables ([N-|A⋃B|, A-B; B-A, |A∩B|], N = number of all genes, A = gene set in each 5xFAD gene lists and B = gene set in each AMP-AD modules), to calculate the p-value of overlap between the 5xFAD gene lists and AMP-AD modules.

#### WGCNA analysis

A matrix filtered by genes with more than 1 TPM and without an outlier sample (both cortex and hippocampus from that sample were removed) was used to do a weighted gene correlation network analysis (WGCNA). Parameters used are: soft power =15, min. module size =50 and MEDissThres = 0.3. We identified significant modules by calculating the correlation with the traits, then we proceeded to plot the behavior per sample of the genes in the blue and dark olive module, by using bar plot and the eigengene profile. Genes from both modules were used for a GO term analysis using Metascape (https://metascape.org).

### Statistics

Every reported *n* is the number of biologically independent replicates. No statistical methods were used to predetermine sample sizes; however, our sample sizes are similar to those reported in recently published similar studies ^9,13^. Behavioral, biochemical, and immunohistological data were analyzed using either Student’s t-test, one-way ANOVA or two-way ANOVA using GraphPad Prism Version 8 (La Jolla, CA). Bonferroni’s and Tukey’s post hoc tests were employed to examine biologically relevant interactions from the two-way ANOVA. *p<0.05, **p<0.01, ***p<0.001 and ***p<0.0001. Statistical trends are accepted at p<0.10 (^#^). Data are presented as raw means and standard error of the mean (SEM).

## Data Records

The results published here are in whole based on data available via the AD Knowledge Portal (https://adknowledgeportal.org). The AD Knowledge Portal is a platform for accessing data, analyses, and tools generated by the Accelerating Medicines Partnership (AMP-AD) Target Discovery Program and other National Institute on Aging (NIA)-supported programs to enable open-science practices and accelerate translational learning. The data, analyses and tools are shared early in the research cycle without a publication embargo on secondary use. Data is available for general research use according to the following requirements for data access and data attribution (https://adknowledgeportal.org/DataAccess/Instructions).

For access to the data see: https://doi.org/10.7303/syn23628482

Data can be accessed in an interactive matter at UCI Mouse Mind Explorer (mouse.mind.uci.edu).

## Technical Validation

### 5xFAD mice show behavior impairment

5xFAD and wild-type littermate mice were aged to 4, 8, 12 and 18 months of age and subjected to a battery of cognitive and behavioral testing tasks, followed by extensive characterization, including long-term potentiation (LTP), immunohistochemistry, biochemistry, and gene expression. Notably, all generated data are explorable in a searchable website (http://mouse.mind.uci.edu/), while raw data (all microscopy images, FASTQ files etc.) are deposited at the AD Knowledge Portal (https://adknowledgeportal.synapse.org/). 5xFAD mice failed to gain weight from 8 months of age, compared to WT mice, and this was most prominent in female mice (Figure 1A and B). Motor impairments were evident in 5xFAD mice at 18 months of age, both by the distance traveled and velocity in the open field test (Figure 1E and G, respectively), with a preference to the center of the arena at 8 months of 5xFAD were observed relative to the WT (Figure 1 C and D). Prominent differences were measured in the elevated plus maze at all timepoints and were present for both male and female 5xFAD mice. 5xFAD spent more time in the open arms, and less time in the closed arms indicating decreased anxiety behaviors (Figure 1I-L) (in contrast to no differences noted in open field). Of note, we have previously shown similar changes in EPM performance in a mouse model of selective hippocampal neuronal loss^14^. No changes were observed in contextual fear conditioning (Figure 1M-N). Notably, 4 month old 5xFAD mice showed longer latencies to fall on rotarod compared to wild-type mice (Figure 1O), which was driven more so by female mice (Figure 1P), however, reduced motor performance was seen at all subsequent age groups and no genotype differences observed (Figure 1O).

**Figure 1.**
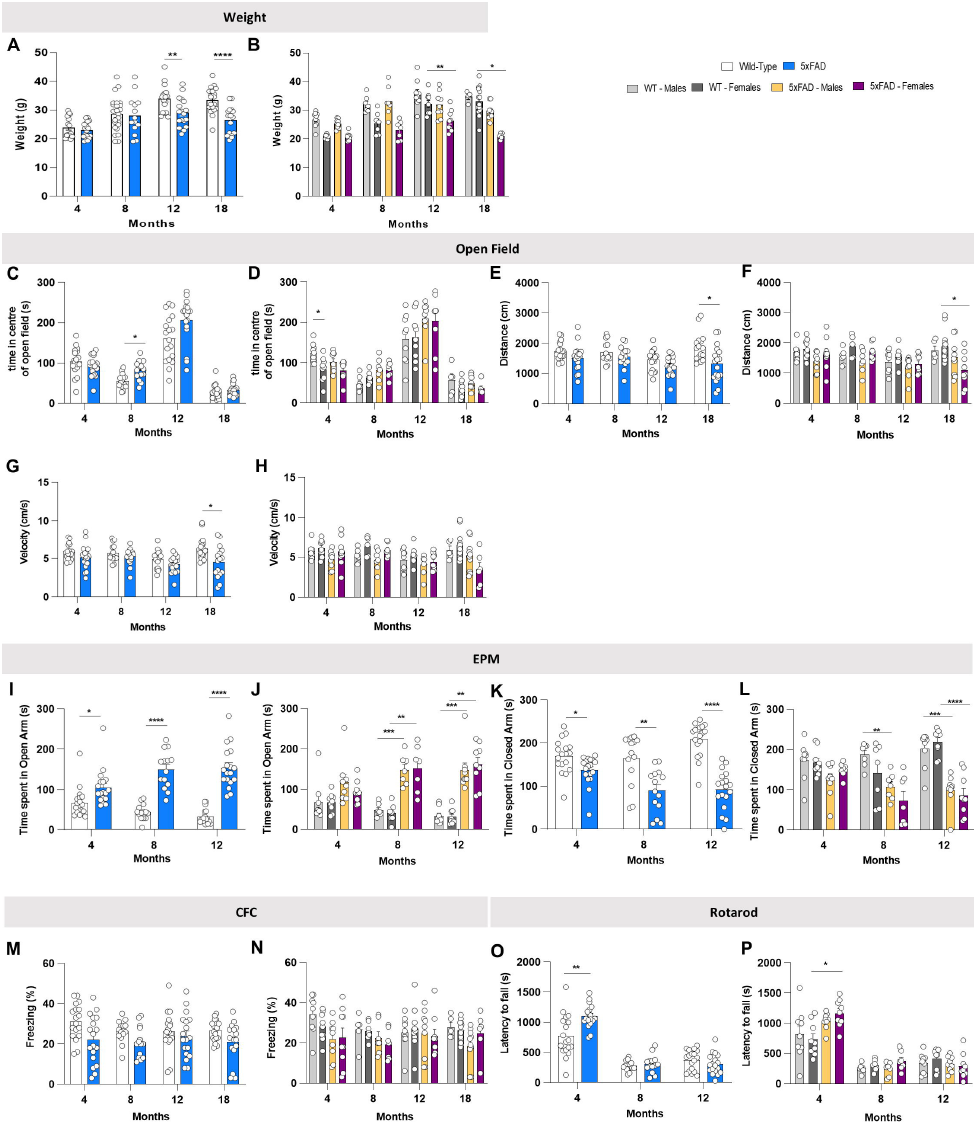
Behavioral tasks reveal age-related changes in both WT and 5xFAD mice. *A-B)* 5xFAD at 12- and 18-month of age show less weight gain than their littermate WT; this effect is higher on females. *C-H)* The open field test reveals deficits in distance traveled and velocity at 18 months 5xFAD (*E and G, respectively*). *I-L)* 4, 8 and 12-month old 5xFAD mice spend more time in the open arms and less time in the closed arms of the elevated plus maze. *M-N)* There is no effect of either age nor genotype on the contextual fear conditioning. *O-P)* On the rotarod, 4-month-old 5xFAD time of latency is higher than WT, the effect being more on females. Data are represented as mean ± SEM. *P ≤ 0.05, **P ≤ 0.01, *** P ≤ 0.001, **** P ≤ 0.0001, n = 9-10 per group.32eo1pgirwhu342o58oy374=

### 5xFAD mice display impaired LTP and synaptic transmission

We assessed short- and long-term synaptic plasticity using acute hippocampal slice preparation from WT and 5xFAD mice. Field EPSPs were evoked in the proximal apical dendrites in field CA1b during stimulation of Schaffer-commissural projections in CA1a and LTP was induced using theta burst stimulation. Across all ages, 4, 8 and 12 months, we found that theta bust-induced LTP produced significant reductions in the level of potentiation 50-60 min post-induction. Beginning at 4 months (Figure 2A,B), potentiation was reduced in both male and female 5xFAD mice compared to WT mice. LTP remained impaired in both sexes in slices from 8 and 12 months 5xFAD mice relative to WT controls (Figure 2C-F). Baseline synaptic transmission was also evaluated for all ages and revealed that fEPSP responses in slices from 12 months 5xFAD mice were markedly reduced compared to WT slices, and furthermore, the decrease in field responses was observed in both sexes in 5xFAD mice relative to controls (Figure 2G). Evaluating changes in paired-pulse facilitation showed that at 12 months of age frequency facilitation was significantly reduced in slices from 5xFAD mice compared to WT controls (Figure 2H, top panel), which is due to the difference observed in the males relative to their controls (Figure 2H, bottom panel). No differences were observed in paired-pulse facilitation in slices from female 5xFAD and WT mice at 12 months of age (Figure 2H, bottom panel). Altogether, these synaptic data suggest deficits in LTP and synaptic transmission in 5xFAD mice beginning at 4 months, and worsening with age.

**Figure 2.**
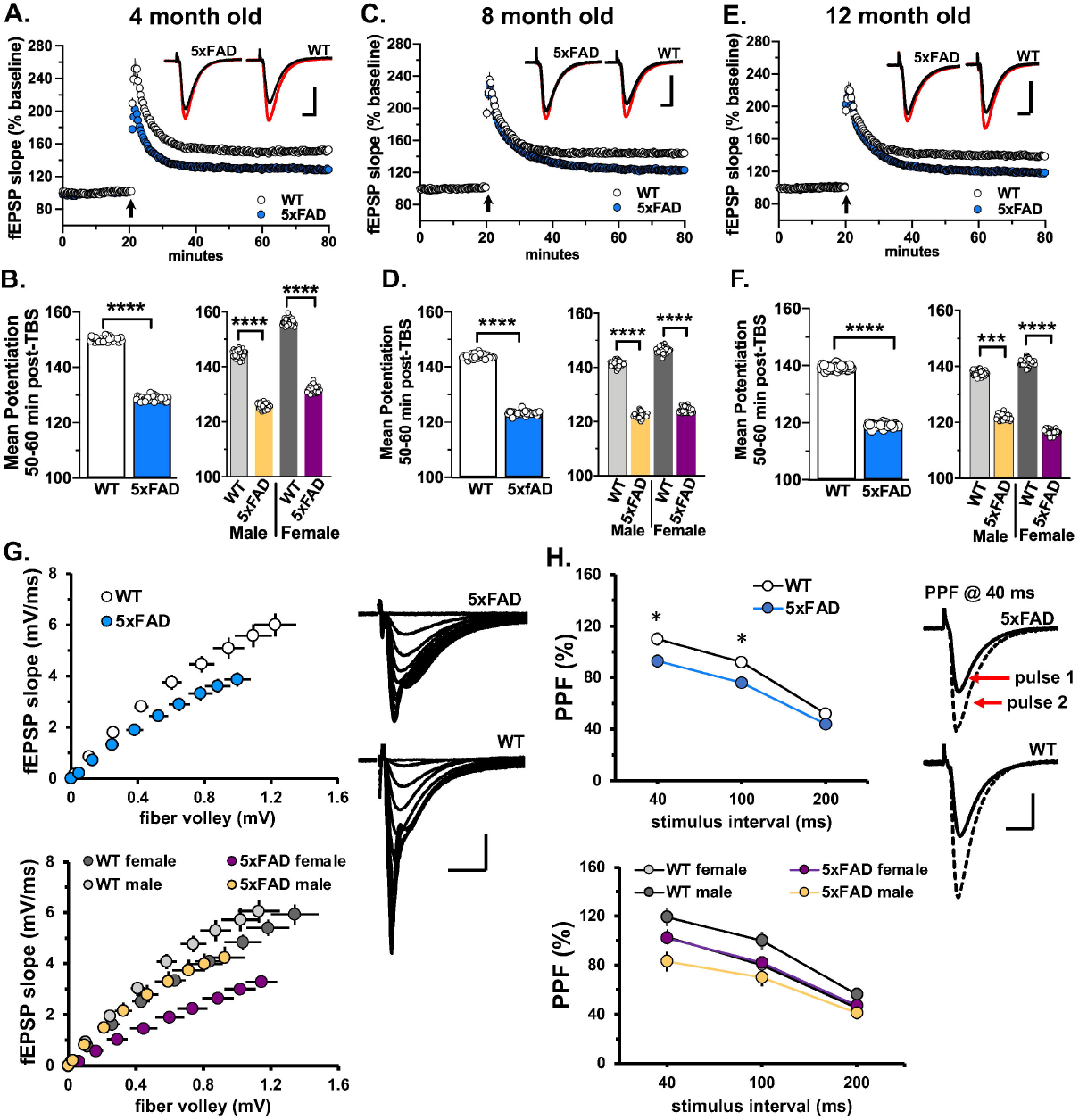
5xFAD Mouse Model Produces LTP Impairments. Theta burst-induced LTP is impaired in 5xFAD mouse model. Hippocampal slices were prepared from 4, 8, and 12-month-old male and female WT and 5xFAD mice. *A)* Time course for theta-burst induced (black arrow) LTP shows that the level of potentiation is notably reduced in slices from 4-month-old 5xFAD mice relative to slices from WT controls. Insets show field synaptic responses collected during baseline (black line) and 1 hour after theta burst stimulation (red line). Scale: 1 mV/ 5 ms. *B)* Left bar graph, Group summary of mean potentiation (± SEM) during the last 10 min of recording in slices from 4 months WT and 5xFAD mice (F1,35 = 35.8, p<0.0001). Right bar graph, Mean potentiation in slices from 4 months male and female WT and 5xFAD (male, F1,17 = 19.9, p = 0.003; female, F1,16 = 23.0, p = 0.0002). *C)* Time course for theta-burst induced LTP shows that the level of potentiation is reduced in slices from 8 months old 5xFAD mice relative to WT controls. Insets show field synaptic responses collected during baseline (black line) and 1 hour after theta burst stimulation (red line). Scale: 1 mV/ 5 ms. *D)* Left bar graph, Group summary of mean potentiation collected during the last 10 min of recording in slices from 8 months WT and 5xFAD mice (F1,38 = 64.2, p < 0.0001). Right bar graph, Mean potentiation in slices from 8 months male and female WT and 5xFAD (males, F1,19 = 31.6, p < 0.0001; females, F1,17 = 32.8, p < 0.0001). (E) Time course for theta-burst induced LTP again shows that the level of potentiation is markedly lower in slices from 12 month old 5xFAD mice relative to WT controls. Insets show field synaptic responses collected during baseline (black line) and 1 hour after theta burst stimulation (red line). Scale: 1 mV/ 5 ms *F)* Left bar graph, Group summary of mean potentiation during the last 10 min of recording in slices from 12 months WT and 5xFAD mice (F1,36 = 64.4, p < 0.0001). Right bar graph, Mean potentiation in slices from 12 months male and female WT and 5xFAD (male, F1,17 = 16.7, p = 0.0008; female, F1,17 = 59.3, p < 0.0001). *G)* The input/output curve measuring the amplitude of the fiber volley relative to the fEPSP slope at 12 months was significantly different between WT and 5xFAD group (top panel, F 1,36 = 22.8, p < 0.0001), and gender (bottom panel, male, F 1,17 = 4.5, p = 0.049; female, F 1,17 = 34.4, p < 0.0001). Field traces on the right show representative synaptic responses collected during generation of an input/output curve in a slice from a 12-month-old WT and 5xFAD mouse. Scale: 1 mV/ 5 ms. (H) Paired-pulse facilitation was measured at 40, 100, and 200 ms intervals. Top panel, At 12 months of age, PPF is significantly reduced in slices from 5xFAD mice with respect to age-matched WT controls (F 1,36 = 5.8, p = 0.02). Bottom panel. This effect is due to the notable separation in PPF at 40 and 100 ms stimulus intervals between male 5xFAD and WT controls (males, F 1,17 = 9.6, p = 0.006; females, F 1,17 = 0.03, p = 0.86). Field traces on the right represent a pair of evoked responses at 40 ms collected in a slice from a 12 months male 5xFAD and WT mouse. Scale: 1 mV/ 5 ms.

### Age-related increases in Aβ plaque accumulation in 5xFAD mice

Immunofluorescence was performed on every 12^th^ section throughout the rostral-caudal axis of the brain. All images are available for exploration and download at AD Knowledge Portal (https://adknowledgeportal.synapse.org/). 5xFAD males and females were stained with Thio-S for characterization of fibrillar amyloid plaques at 4-, 8-, 12- and 18-month timepoints. Absence of plaque pathology was evident throughout the entire brain in WT but was present and exacerbated by age in the 5xFAD as expected (Figure 3A). Plaque pathology was noticeable throughout the rostral-caudal axis of the brain by 4 months of age (Figure 3B). Notably, the initial plaques that develop by 4 months of age are typically compact and circular, but over time appear more irregular and develop a diffuse halo in the subiculum, CA1 and cortex (12-18 months of age; Figure 3C). Importantly, this halo effect is similar to what is observed in the human brain (data not shown).

**Figure 3.**
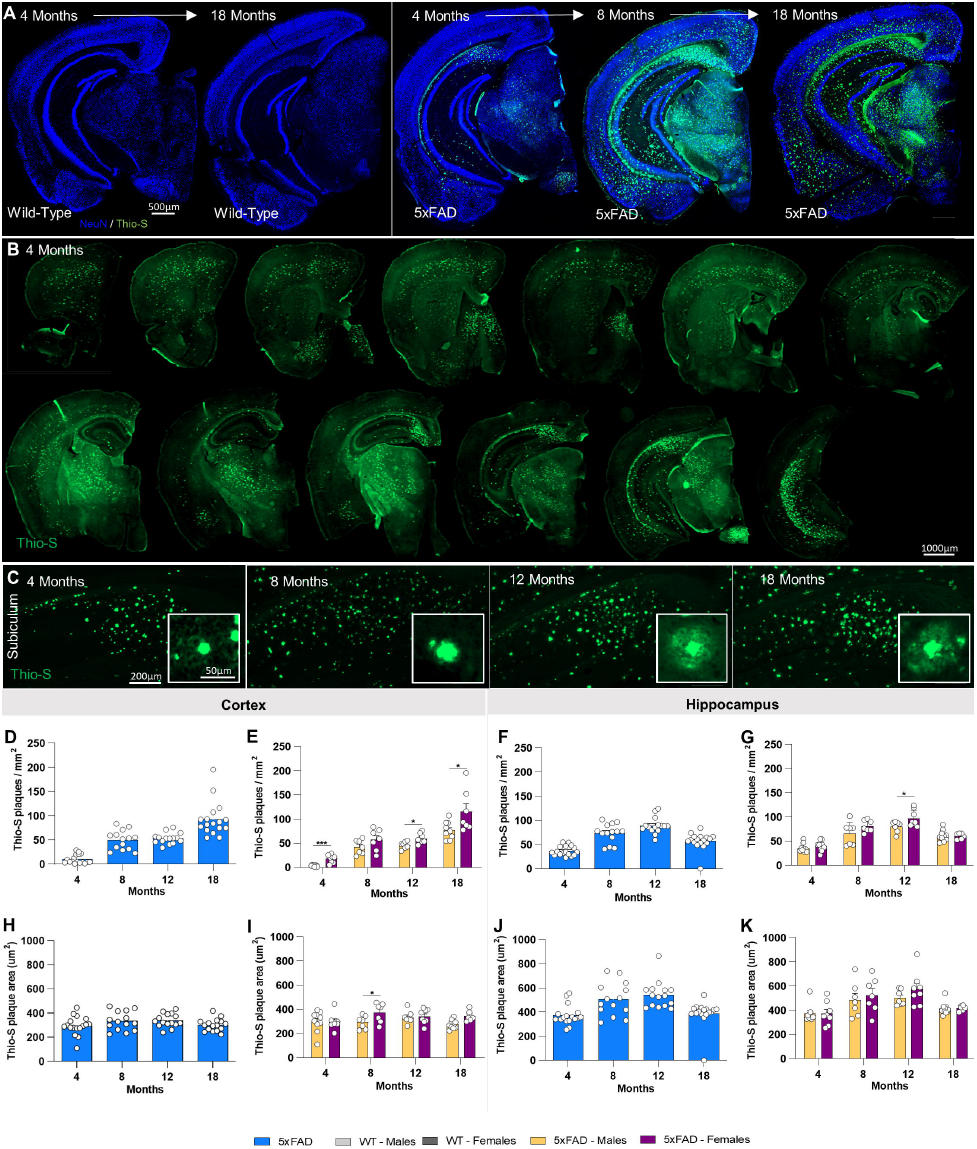
Fibrillar amyloid plaques increase in size and number in 5xFAD aged mice. 5xFAD plaque burden was assessed with Thio-S staining at each time point. *A-B)* Representative stitched brain hemispheres of WT and 5xFAD shown with Thio-S staining at the 4- and 18-month and 4, 8, and 18 mo timepoints respectively, counter stained for NeuN. *B)* Representative stitched whole brain hemispheres of 5xFAD (rostral to caudal) shown with Thio-S staining at the 4 month timepoint. *C)* Representative images of plaques in 5xFAD mice across timepoints displaying a “halo” effect at 12 and 18 months. *D-G)* Quantification for number of Thio-S positive plaques in the cortex and hippocampus by genotype and sex. *H-K)* Quantification of average plaque area in the cortex and hippocampus by genotype and sex. Data are represented as mean ± SEM. *P ≤ 0.05, **P ≤ 0.01, *** P ≤ 0.001, **** P ≤ 0.0001, n = 6 per sex per age.

Absolute values with time are not necessarily a reflection of pathology since they were processed at different time, but relationships within a given time point are valid. As expected, plaque number increased in both the cortex and hippocampus of males and females between 4 and 8 month and with additional increases in the cortex by 18 months (Figure 3D and F). Clear sex differences were seen at 4, 12 and 18 months of age with female 5xFAD mice having a higher number of plaques in the cortex than male 5xFAD (Figure 3E). Plaque size increased with age in the hippocampus, followed by an overall reduction between 12 and 18 months of age, likely reflecting increased plaque compaction (Figure 3J), while cortical plaque size remained stable across the lifespan (Figure 3H). No prominent sex differences were seen in plaque size (Figure 3I and K).

To supplement quantification of plaque load, measurements of Aβ40, and Aβ42, from microdissected hippocampus and cortex, were performed in detergent soluble and insoluble fractions. Prominent increases in soluble Aβ40 and Aβ42 levels were seen at 18 months in both regions (Figure 4A-D; 4E-H). In concordance with plaque numbers, insoluble Aβ is elevated in the cortex in an age dependent fashion (Figure 4I-L), while the hippocampus shows a plateau from 8 months of age (Figure 4M-P), consistent with plaque numbers. Again, female mice tend to have higher levels of insoluble Aβ, with significance for Aβ40 seen at 12 months of age (Figure 4J and N). Plasma Aβ40 and Aβ42 levels are elevated from 8 months of age with Aβ42 levels higher at 8 and 12 months than Aβ40, with no differences between sexes (Figure 4Q and R).

**Figure 4.**
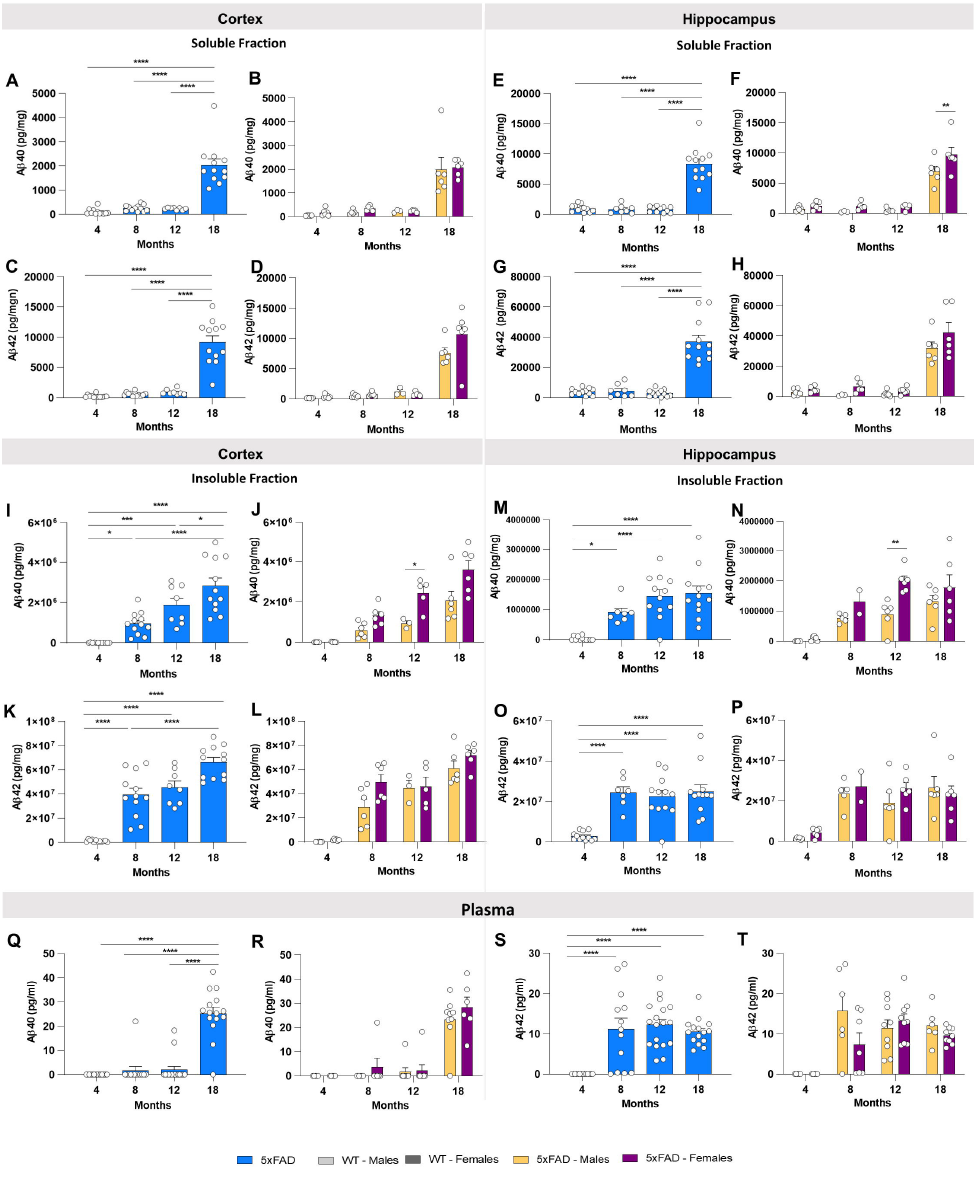
Protein differences observed with age and in 5xFAD mice. Levels of Aβ were quantified in microdissected hippocampi and cortices via Mesoscale Multiplex technology. *A-H)* Levels of Aβ40 and Aβ42 were measured in the soluble fraction of cortex and hippocampus, respectively, with age-related increases in the level of Aβ40 and Aβ42 shown in cortex and hippocampus of 5xFAD mice. *I-P)* Increases in levels of insoluble Aβ40 and Aβ42 were seen in cortex and hippocampus with age. *Q-T)* Increases of plasma levels of Aβ40 at 18 months of age 5xFAD and Aβ42 at 8-, 12- and 18-month old 5xFAD. Data are represented as mean ± SEM. *P ≤ 0.05, **P ≤ 0.01, *** P ≤ 0.001, **** P ≤ 0.0001, n = 6 per group.

### Age-related microgliosis in 5xFAD mice

Immunostaining for the microglial marker IBA1 revealed increases in microglial densities from 8 months of age in the cortex of 5xFAD mice, and from 4 months of age in the hippocampus (Figure 5A and B). Microglia clustered around dense core plaques, as expected. Microglial numbers remained stable in WT mice across the lifespan but increased in 5xFAD mice (Figure 5C and E), mirroring the plaque load. Concordantly, female 5xFAD mice tend to have increased microglial densities, while no sex differences are observed in WT mice (Figure 5D and F).

**Figure 5.**
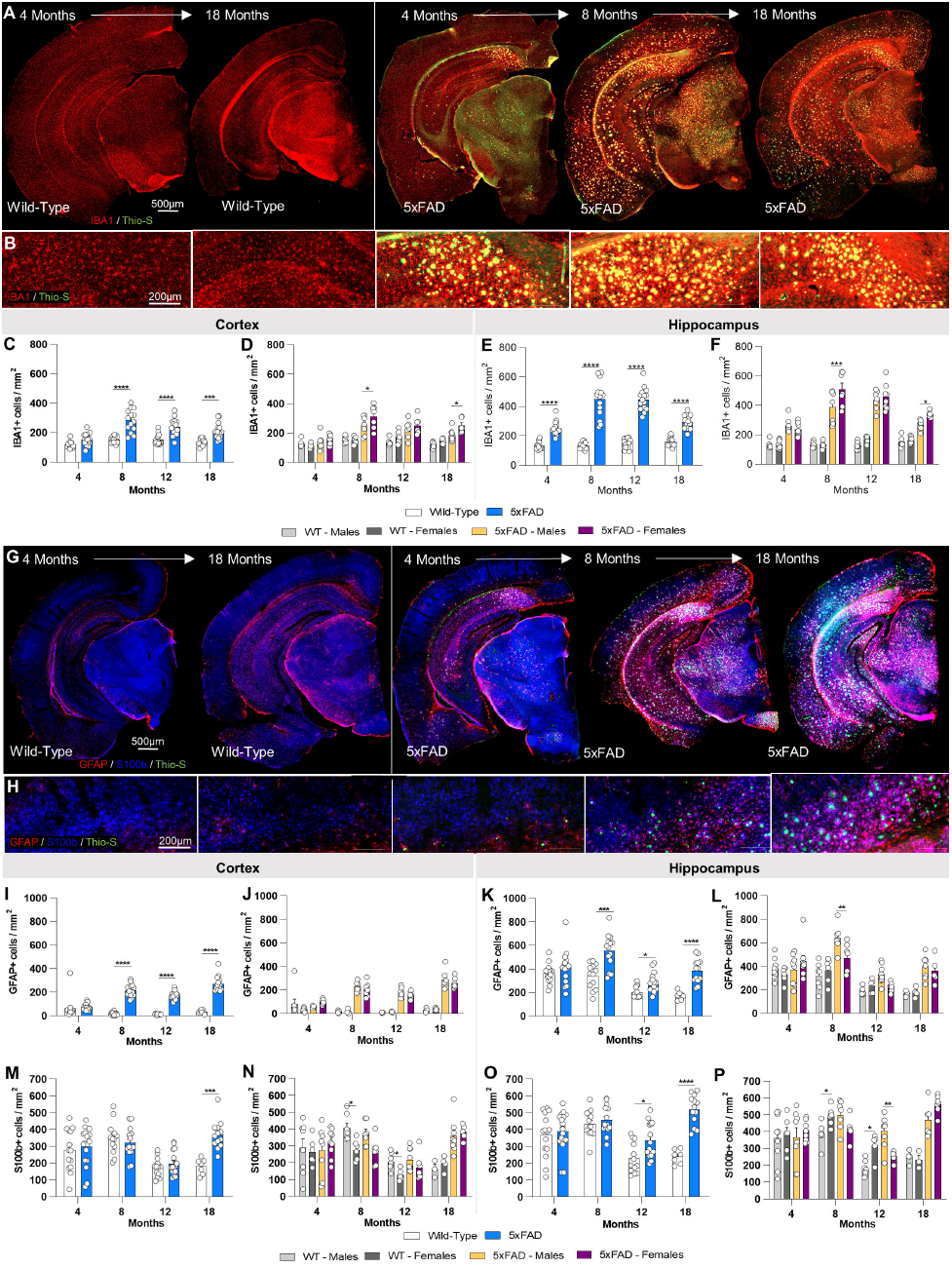
Immunostaining of microglia and astrocytes. Brains of mice at each timepoint were sliced and immunostained for IBA1, GFAP and S100ß to reveal any changes in microglial, astrocytic. *A-B)* Representative stitched brain hemispheres of WT and 5xFAD shown with IBA1/Thio-S staining at the 4- and 18-month and 4, 8, and 18 months timepoints, respectively. *C-F)* IBA1 immunostaining for microglia reveals both age-related changes in WT and 5xFAD microglial number, and differences between genotypes in cortex and hippocampus. *G-H)* Representative stitched brain hemispheres of WT and 5xFAD shown with GFAP/ S100ß/Thio-S staining at the 4- and 18-month and 4, 8, and 18 months timepoints, respectively. *I-P)* Astrocyte number is assessed via GFAP (I-L)) and S100ß staining (*M-P*) in the cortex and hippocampus. Data are represented as mean ± SEM. *P ≤ 0.05, **P ≤ 0.01, *** P ≤ 0.001, **** P ≤ 0.0001, n = 6 per group.

### Age-dependent astrocyte reactivity in 5xFAD mice

To quantify astrocyte numbers and reactivity state, IHC for S100b and GFAP was performed (Figure 5G and H). S100b is a nuclear transcription factor expressed by all astrocytes, while GFAP is expressed by hippocampal astrocytes, but in the cortex is only expressed by “reactive” astrocytes. Immunostaining for S100b shows significantly increased astrocyte densities at 18 months of age in 5xFAD mice compared to WT mice in the cortex, and from 12 months of age in the hippocampus (Figure 5M-P). GFAP+ astrocytes mirror S100b trends in the hippocampus, with elevated GFAP+ cells seen from 8-18 months of age (Figure 5K and L). Astrocytes in the cortex are observed to switch on GFAP expression in the vicinity of plaques (Figure 5H), and GFAP+ astrocyte numbers hence follow plaque numbers (Figure 5I and J).

### Age dependent dystrophic neurite accumulation in 5xFAD mice

Dense core plaques are surrounded by dystrophic neurites, which can be observed via immunostaining for the lysosome-associated membrane protein 1 (LAMP-1). LAMP1 and Thio-S staining was performed in all timepoints of WT and 5xFAD mice (Figure 6A and B). We quantified both Thio-S and LAMP1 staining as a % load (i.e., brain area covered by the positive signal); Thio-S increased in an age dependent fashion, with a much higher load in the hippocampus compared to the cortex (Figure 6C, D, I, J) consistent with the plaque number quantified in Figure 3. LAMP1 load increases with plaque load (Figure 6E, F, K, L) but reached a plateau at 8 and 12 months of age in cortex and hippocampus respectively, suggesting that while both plaque load and dystrophic neurites increase with age, the associated halo of dystrophic neurites does not increase proportionally. As such, the ratio between Thio-S and LAMP1 load reduces with age (Figure 6G, H, M, N).

**Figure 6.**
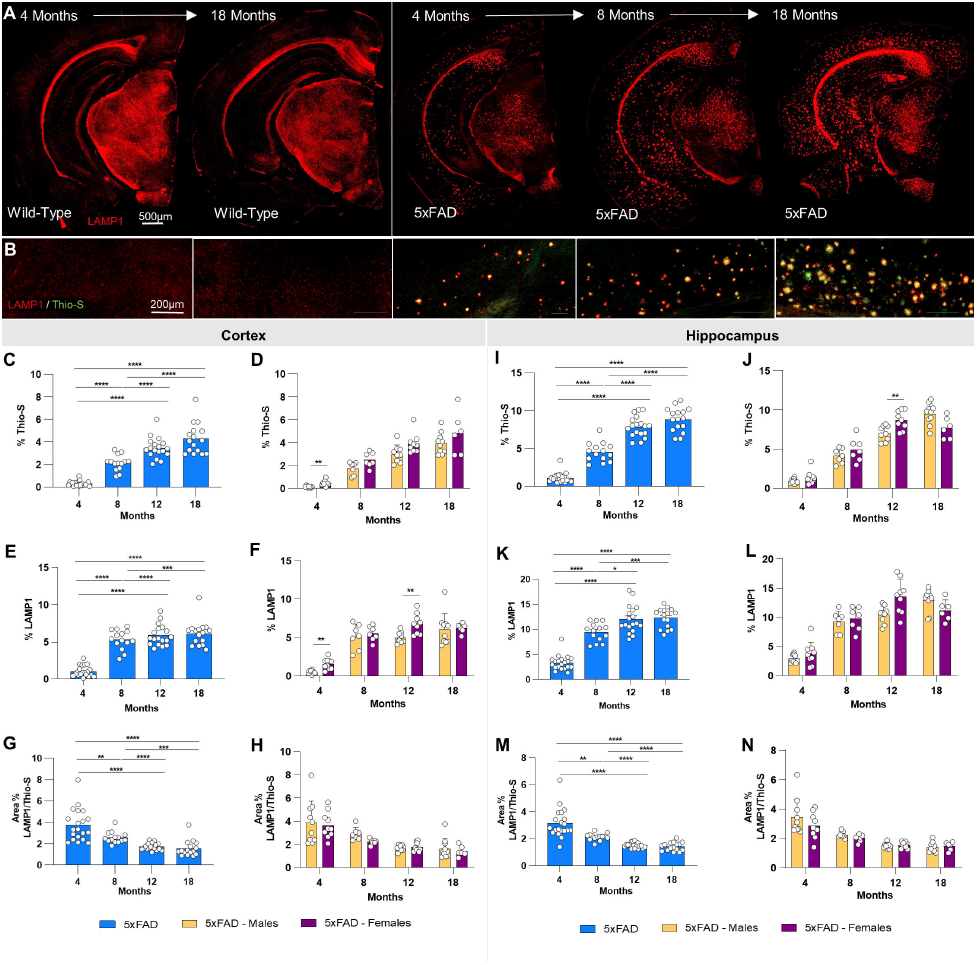
Immunostaining of lysosomes. *A-B)* Representative stitched brain hemispheres of WT and 5xFAD shown with LAMP1/Thio-S staining at 4 and 18 months, and 4-, 8- and 18-months timepoints, respectively. *C-D, I-J)* Quantification of Thio-S in cortex and hippocampus. *E-F, K-L*) LAMP1 immunostaining for lysosomes reveals age-related changes of 5xFAD mice in percent area of the cortex and hippocampus covered by LAMP1. *G-H)*. In quantifying the ratio of LAMP1/Thio-S coverage, there was an age-related decrease, but no sex-related changes in the cortex. *M-N)* A ratio of the percent area coverages of LAMP1 and Thio-S reveals age-related changes in the hippocampus of 5xFAD mice and no sex-related changes.

### Gene expression changes in 5xFAD mice

Differentially expressed genes (DEG’s) were calculated for comparisons between WT and 5xFAD mice for both the cortex and hippocampus at each timepoint. These data are explorable at http://mouse.mind.uci.edu and at https://adknowledgeportal.synapse.org/. The number of DEG’s was higher in the hippocampus at each timepoint than cortex and increased with age in both brain regions (Figure 7A). Notably, 18-month 5xFAD mice showed a large increase in upregulated DEG’s in both brain regions, when downregulated genes were also observed. To evaluate overlap in DEG’s between brain regions and across the lifespan of 5xFAD mice we produced a chart (Figure 7B) highlighting downregulated genes (blue) and upregulated genes (red). Substantial overlap was seen in the upregulated genes between hippocampus and cortex, though a set of unique upregulated genes seen in the hippocampus at 18 months (Figure 7G). Overall, far fewer downregulated DEG’s were seen, but a substantial unique set of genes materialized at 18 months in the hippocampus (Figure 7I). Gene ontology of common upregulated genes (upregulated in 4 out of 4 of the timepoints for hippocampus) identified pathways involved in inflammation, as expected (Figure 7C, D), while common downregulated genes (in at least 2 out of 4 of the timepoints for hippocampus) related with pathways associated with synaptic transmission and signaling (Figure 7E, F). Gene ontology analyses of the unique DEG’s at 18 months in the hippocampus revealed pathways associated with vascular development for upregulated genes (Figure 7G, H), and synaptic transmission for the downregulated genes (Figure 7I, J).

**Figure 7.**
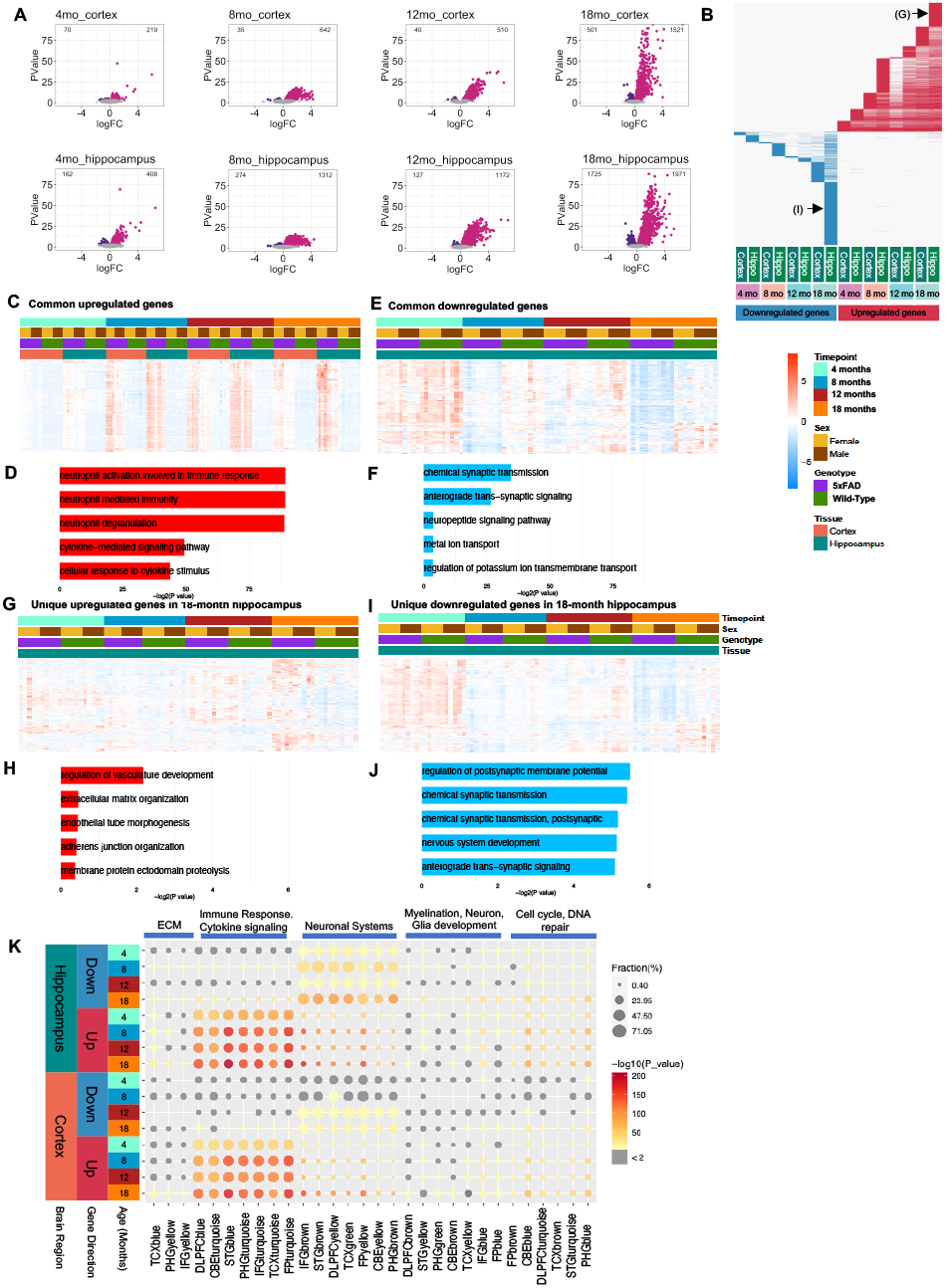
Differential gene expression analysis of the 5xFAD time course. A) Comparisons of 5xFAD and WT were done across different timepoints and tissues. Upregulated genes are labeled in pink and down regulated genes are labeled in purple. Number of differential expressed genes is displayed in the upper corners of the volcano plot. Parameters FDR < 0.05. B) Comparison of differential expressed genes across timepoint and tissue. Upregulated genes in red, downregulated genes in blue. Each column represents a set of genes for a different time point, each row represents each one of the differentially expressed genes. Unique upregulated and downregulated gene sets representing in Figure 7G and 7I are also indicated as (G) and (I) in this panel. C-D) Heatmap and GO Term analysis for common genes upregulated. E-F) Common downregulated genes, G-H) Unique genes upregulated at 18 months in hippocampus, I-J) Unique genes downregulated at 18 months in hippocampus. K) Comparison of differentially expressed genes against AMP-AD modules. Size of the dot represents the fraction and color represents how much this fraction is significant.

To understand the relevance of these gene expression changes to human AD, we compared these DEG’s to identified AMP-AD modules reflecting gene expression changes in human AD samples^15^. Significant overlap was seen in both down- and up-regulated genes, with the strongest overlap seen in the 5xFAD hippocampus at 18 months of age (Figure 7K).

To further understand gene expression in 5xFAD mice in the context of networks we performed WGCNA to recover 11 modules, which we correlated with genotype, age, and previously described phenotypic characterization (Figure 8A). We found that the Blue module (681 genes) is positively correlated with the 5xFAD genotype (P-value = 4e-23), while the DarkOliveGreen module (524 genes) is negatively correlated with the 5xFAD genotype (P-value = 0.09). These modules are also correlated with different phenotypes and some specific gene expression levels. For example, the Blue module is strongly positively correlated with microglia count (P-value = 6e-29), plaque count (P-value = 5e-23), among other phenotypes. Overall, genes in the Blue module (Figure 8B) increase expression in 5xFAD with age, whereas genes in the DarkOliveGreen module (Figure 8C) decrease expression in 5xFAD with age. GO term analysis of genes in the Blue module reveals that this module is enriched in genes involved in immune systems response (Figure 8D) that are primarily expected to be microglial, although a few astrocytic genes such as GFAP are also in this module. By contrast, GO terms for the DarkOliveGreen Module are primarily neuronal in nature (Figure 8E). Overall, distinct gene modules correlate with phenotypic changes in our 5xFAD dataset.

**Figure 8.**
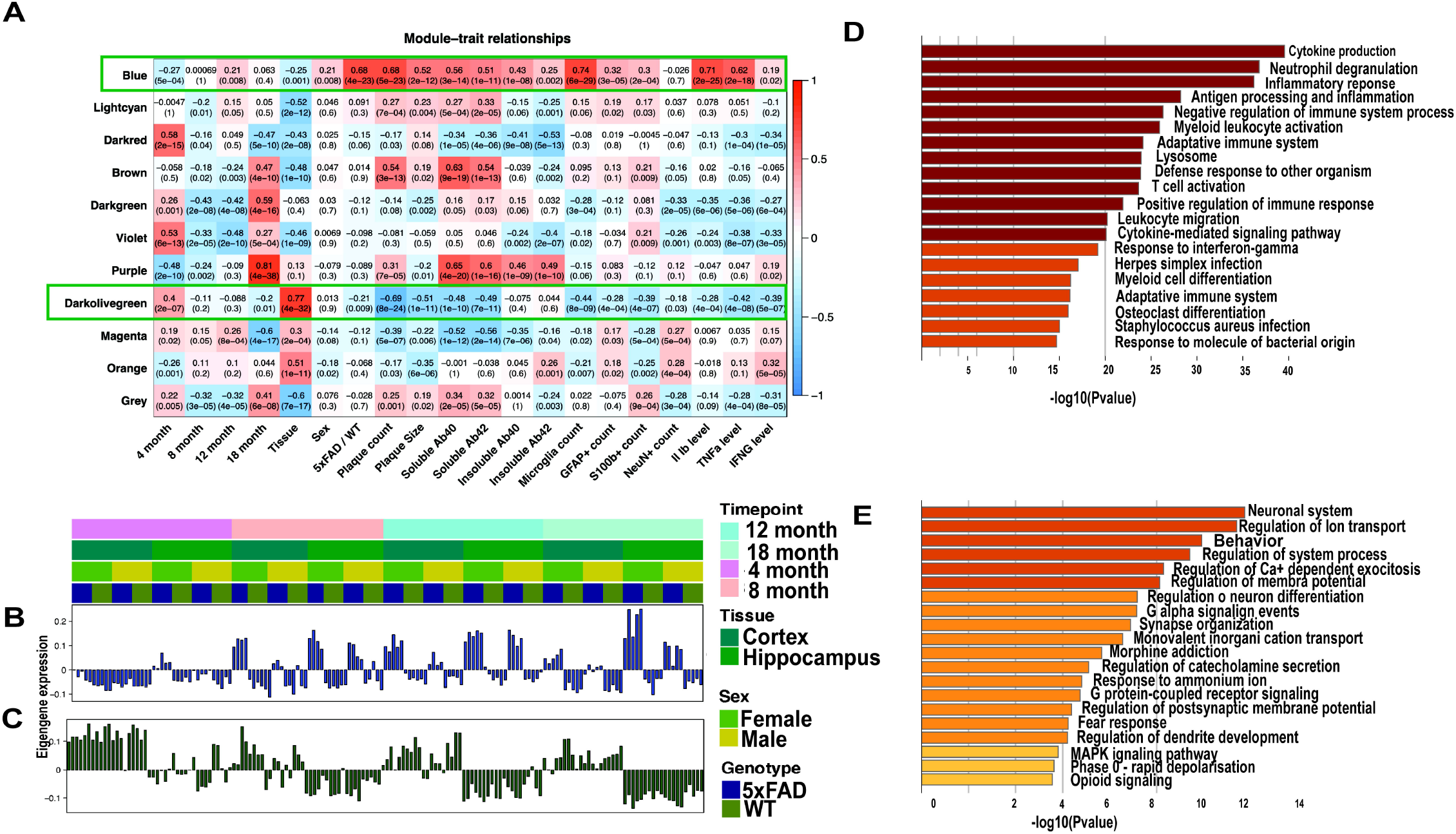
Gene expression during progression of the 5xFAD phenotype. A) Matrix with the Module-Trait Relationships (MTRs) and corresponding p-values between the detected modules on the y-axis and selected AD traits on the x-axis. The MTRs are colored based on their correlation: red is a strong positive correlation, while blue is a strong negative correlation. B) Bar plots for the eigengene expression and heatmap of the genes in the blue module. C) Bar plots for the eigen expression of the genes in the dark olive-green module. D) and E) Gene ontology analysis for genes of the blue and dark olive-green module respectively.

### Usage Notes

A critical goal of the research community is to develop and characterize animal models of Alzheimer’s disease that represent the various stages and pathologies that define the human disease. These models are important for the cross sectional understanding of the aging-related changes that lead to the development of AD, which is not easily achieved using human brain samples that represent the end stage (and/or one time point) of the disease, and in order to develop and test therapeutics^16^ with high translational value. The identification of risk-associated polymorphisms to late-onset AD over the past several decades is aiding our understanding of the disease, and directing new therapeutic avenues, for example against microglia^13,17-20^. Given the pronounced differences between humans and mice, modeling this complex disease of aging has proven challenging, with salient differences in lifespan, and in the sequences and processing of the key proteins that define the prominent pathologies of the AD brain (such as plaques (APP) and tangles (tau)). As such, it is unlikely that a single animal/mouse model will recapitulate all the pathologies seen in the human brain, and thus multiple animal models will be needed to model different aspects of the disease. Furthermore, given the age-related and progressive nature of the disease it is likely that within any animal model the appropriate ages will need to be defined. Many existing mouse models of AD have utilized human APP alongside familial/early onset mutations to drive amyloidogenesis and recapitulate plaque pathology and have been useful for developing therapies that can mitigate this aspect of the disease such as via Aβ immunotherapy^21-24^. One of the most widely utilized mouse models by the AD research community is the 5xFAD mouse – here we sought to phenotype and characterize the 5xFAD mice model at 4, 8, 12 and 18 months of age within the MODEL-AD Consortium. We provide in depth phenotyping data that reaffirm that this model develops robust amyloid pathology ^4^, and downstream microgliosis and inflammation ^25-27^, reactive astrocytes, and the induction of dystrophic neurites ^28^. We also show robust impairments in long-term potentiation ^29^, and specific deficits in certain behavioral tasks ^4,7^. Plaque pathology is reproducible and develops initially within the subiculum and then spreads throughout the hippocampus and cortex. Notably, we show a sex difference with female mice developing pathology prior to male mice; this is explained by increased expression of the Thy1 promoter used to drive the transgenes in this model which has an estrogen response element ^30,31^ resulting in generation of higher levels of Aβ ^4,32^. Furthermore, we provide gene expression data from all timepoints, and find that upregulated genes mostly represent the inflammatory response of the glia to the Aβ plaques while downregulated genes are associated with synaptic and neuronal function. Critically, we show that different brain regions (i.e. cortex and hippocampus) have both common and unique gene expression responses to the pathology, and that these changes better recapitulate the human AD brain with increased age, with 18 months 5xFAD mice showing the most concordance. All data are explorable in an interactive fashion at http://mouse.mind.uci.edu, while raw data can be downloaded at the AD Knowledge Portal (https://adknowledgeportal.org), including histology from the entire rostral-caudal axis showing the spatial and temporal progression of pathology. The MODEL-AD consortium is developing and characterizing new animal models based on GWAS identified AD risk variants, humanization of key genes, and diverse genetic backgrounds and these data and the mice will be available in a similar fashion to allow researchers to explore and select the appropriate animal model and age for their needs. Existing models such as the 5xFAD mice have value as a robust and consistent model of amyloidosis and the effects of this on the brain, as a model to compare and contrast to new models. Use of standardized protocols of characterization with longitudinal analysis across the lifespan in both sexes should accelerate progress toward targeted therapeutics that will translate with higher efficacy in the clinic.

## Acknowledgements

The animal models in this study were whole or in part created by the Model Organism Development and Evaluation for Late-onset Alzheimer’s Disease (MODEL-AD) consortium funded by the National Institute on Aging. Relevant study strains and characterization data were generated by: the Indiana University/The Jackson Laboratory MODEL-AD Center U54 AG054345 led by Bruce T. Lamb, Gregory W. Carter, Gareth R. Howell, and Paul R. Territo; the University of California, Irvine MODEL-AD Center U54 AG054349 led by Frank M. LaFerla and Andrea J. Tenner. These resources were enhanced by: RF1 AG055104 to Michael Sasner, Gregory W. Carter, and Gareth R. Howell and U54 AG054349 S1-9 to Grant MacGregor, Kim Green, Andre Obenaus, Ian Smith, Xiangmin Xu, and Katrine Whiteson.

## Author Contributions

SF, SK, EAK, GBG, EAK, DPM, JP, DIJ, KMT, EH, CC, NR, JAA, DBV and CJ performed experiments. SF, SK and KNG analyzed the data and wrote the manuscript with contributions from all authors. JN, MW, GRM, AM, AJT, FML, KNG provided advice on the study design, managed the project, and contributed to the manuscript.

## Figure Legends

**Supplemental figure. Phenotyping pipeline of the 5xFAD mouse model.** The process order by which the animals and sample tissue go through within the MODEL-AD phenotyping pipeline at UCI, including behavior, LTP, RNA-seq, histology and biochemical assays.

## References

1 King, A. The search for better animal models of Alzheimer’s disease. Nature 559, S13–S15, doi:10.1038/d41586-018-05722-9 (2018).

2 Drummond, E. & Wisniewski, T. Alzheimer’s disease: experimental models and reality. Acta Neuropathol 133, 155–175, doi:10.1007/s00401-016-1662-x (2017).

3 LaFerla, F. M. & Green, K. N. Animal models of Alzheimer disease. Cold Spring Harb Perspect Med 2, doi:10.1101/cshperspect.a006320 (2012).

4 Oakley, H. et al. Intraneuronal beta-amyloid aggregates, neurodegeneration, and neuron loss in transgenic mice with five familial Alzheimer’s disease mutations: potential factors in amyloid plaque formation. J Neurosci 26, 10129–10140, doi:10.1523/JNEUROSCI.1202-06.2006 (2006).

5 Moechars, D., Lorent, K., De Strooper, B., Dewachter, I. & Van Leuven, F. Expression in brain of amyloid precursor protein mutated in the alpha-secretase site causes disturbed behavior, neuronal degeneration and premature death in transgenic mice. EMBO J 15, 1265–1274 (1996).

6 Vidal, M., Morris, R., Grosveld, F. & Spanopoulou, E. Tissue-specific control elements of the Thy-1 gene. EMBO J 9, 833–840 (1990).

7 Jawhar, S., Trawicka, A., Jenneckens, C., Bayer, T. A. & Wirths, O. Motor deficits, neuron loss, and reduced anxiety coinciding with axonal degeneration and intraneuronal Abeta aggregation in the 5XFAD mouse model of Alzheimer’s disease. Neurobiol Aging 33, 196 e129–140, doi:10.1016/j.neurobiolaging.2010.05.027 (2012).

8 Elmore, M. R. et al. Colony-stimulating factor 1 receptor signaling is necessary for microglia viability, unmasking a microglia progenitor cell in the adult brain. Neuron 82, 380–397, doi:10.1016/j.neuron.2014.02.040 (2014).

9 Spangenberg, E. E. et al. Eliminating microglia in Alzheimer’s mice prevents neuronal loss without modulating amyloid-beta pathology. Brain 139, 1265–1281, doi:10.1093/brain/aww016 (2016).

10 Bradford, M. M. A rapid and sensitive method for the quantitation of microgram quantities of protein utilizing the principle of protein-dye binding. Anal Biochem 72, 248–254, doi:10.1006/abio.1976.9999 (1976).

11 Green, K. N., Khashwji, H., Estrada, T. & Laferla, F. M. ST101 induces a novel 17 kDa APP cleavage that precludes Abeta generation in vivo. Ann Neurol 69, 831–844, doi:10.1002/ana.22325 (2011).

12 Robinson, M. D., McCarthy, D. J. & Smyth, G. K. edgeR: a Bioconductor package for differential expression analysis of digital gene expression data. Bioinformatics 26, 139–140, doi:10.1093/bioinformatics/btp616 (2010).

13 Spangenberg, E. et al. Sustained microglial depletion with CSF1R inhibitor impairs parenchymal plaque development in an Alzheimer’s disease model. Nat Commun 10, 3758, doi:10.1038/s41467-019-11674-z (2019).

14 Rice, R. A. et al. Elimination of Microglia Improves Functional Outcomes Following Extensive Neuronal Loss in the Hippocampus. J Neurosci 35, 9977–9989, doi:10.1523/JNEUROSCI.0336-15.2015 (2015).

15 Wan, Y. W. et al. Meta-Analysis of the Alzheimer’s Disease Human Brain Transcriptome and Functional Dissection in Mouse Models. Cell Rep 32, 107908, doi:10.1016/j.celrep.2020.107908 (2020).

16 Oblak, A. L. et al. Model organism development and evaluation for late-onset Alzheimer’s disease: MODEL-AD. Alzheimers Dement (N Y) 6, e12110, doi:10.1002/trc2.12110 (2020).

17 Elmore, M. R. P. et al. Replacement of microglia in the aged brain reverses cognitive, synaptic, and neuronal deficits in mice. Aging Cell 17, e12832, doi:10.1111/acel.12832 (2018).

18 Zhang, Q. et al. Risk prediction of late-onset Alzheimer’s disease implies an oligogenic architecture. Nat Commun 11, 4799, doi:10.1038/s41467-020-18534-1 (2020).

19 Allen, M. et al. Late-onset Alzheimer disease risk variants mark brain regulatory loci. Neurol Genet 1, e15, doi:10.1212/NXG.0000000000000012 (2015).

20 Hernandez, M. X. et al. Prevention of C5aR1 signaling delays microglial inflammatory polarization, favors clearance pathways and suppresses cognitive loss. Mol Neurodegener 12, 66, doi:10.1186/s13024-017-0210-z (2017).

21 Sevigny, J. et al. The antibody aducanumab reduces Abeta plaques in Alzheimer’s disease. Nature 537, 50–56, doi:10.1038/nature19323 (2016).

22 Buttini, M. et al. Beta-amyloid immunotherapy prevents synaptic degeneration in a mouse model of Alzheimer’s disease. J Neurosci 25, 9096–9101, doi:10.1523/JNEUROSCI.1697-05.2005 (2005).

23 Kitazawa, M., Medeiros, R. & Laferla, F. M. Transgenic mouse models of Alzheimer disease: developing a better model as a tool for therapeutic interventions. Curr Pharm Des 18, 1131–1147, doi:10.2174/138161212799315786 (2012).

24 Crehan, H. et al. Effector function of anti-pyroglutamate-3 Abeta antibodies affects cognitive benefit, glial activation and amyloid clearance in Alzheimer’s-like mice. Alzheimers Res Ther 12, 12, doi:10.1186/s13195-019-0579-8 (2020).

25 Salter, M. W. & Stevens, B. Microglia emerge as central players in brain disease. Nat Med 23, 1018–1027, doi:10.1038/nm.4397 (2017).

26 Heneka, M. T. et al. Neuroinflammation in Alzheimer’s disease. Lancet Neurol 14, 388–405, doi:10.1016/S1474-4422(15)70016-5 (2015).

27 Mandrekar-Colucci, S. & Landreth, G. E. Microglia and inflammation in Alzheimer’s disease. CNS Neurol Disord Drug Targets 9, 156–167, doi:10.2174/187152710791012071 (2010).

28 Gowrishankar, S. et al. Massive accumulation of luminal protease-deficient axonal lysosomes at Alzheimer’s disease amyloid plaques. Proc Natl Acad Sci U S A 112, E3699–3708, doi:10.1073/pnas.1510329112 (2015).

29 Kimura, R. & Ohno, M. Impairments in remote memory stabilization precede hippocampal synaptic and cognitive failures in 5XFAD Alzheimer mouse model. Neurobiol Dis 33, 229–235, doi:10.1016/j.nbd.2008.10.006 (2009).

30 Sadleir, K. R., Eimer, W. A., Cole, S. L. & Vassar, R. Abeta reduction in BACE1 heterozygous null 5XFAD mice is associated with transgenic APP level. Mol Neurodegener 10, 1, doi:10.1186/1750-1326-10-1 (2015).

31 Sadleir, K. R., Popovic, J. & Vassar, R. ER stress is not elevated in the 5XFAD mouse model of Alzheimer’s disease. J Biol Chem 293, 18434–18443, doi:10.1074/jbc.RA118.005769 (2018).

32 Maarouf, C. L. et al. Molecular Differences and Similarities Between Alzheimer’s Disease and the 5XFAD Transgenic Mouse Model of Amyloidosis. Biochem Insights 6, 1–10, doi:10.4137/BCI.S13025 (2013).

