## Supplementary figures and images for "Systematic Phenotyping and Characterization of the 5xFAD mouse model of Alzheimer’s Disease"

### Supplemental figure 1

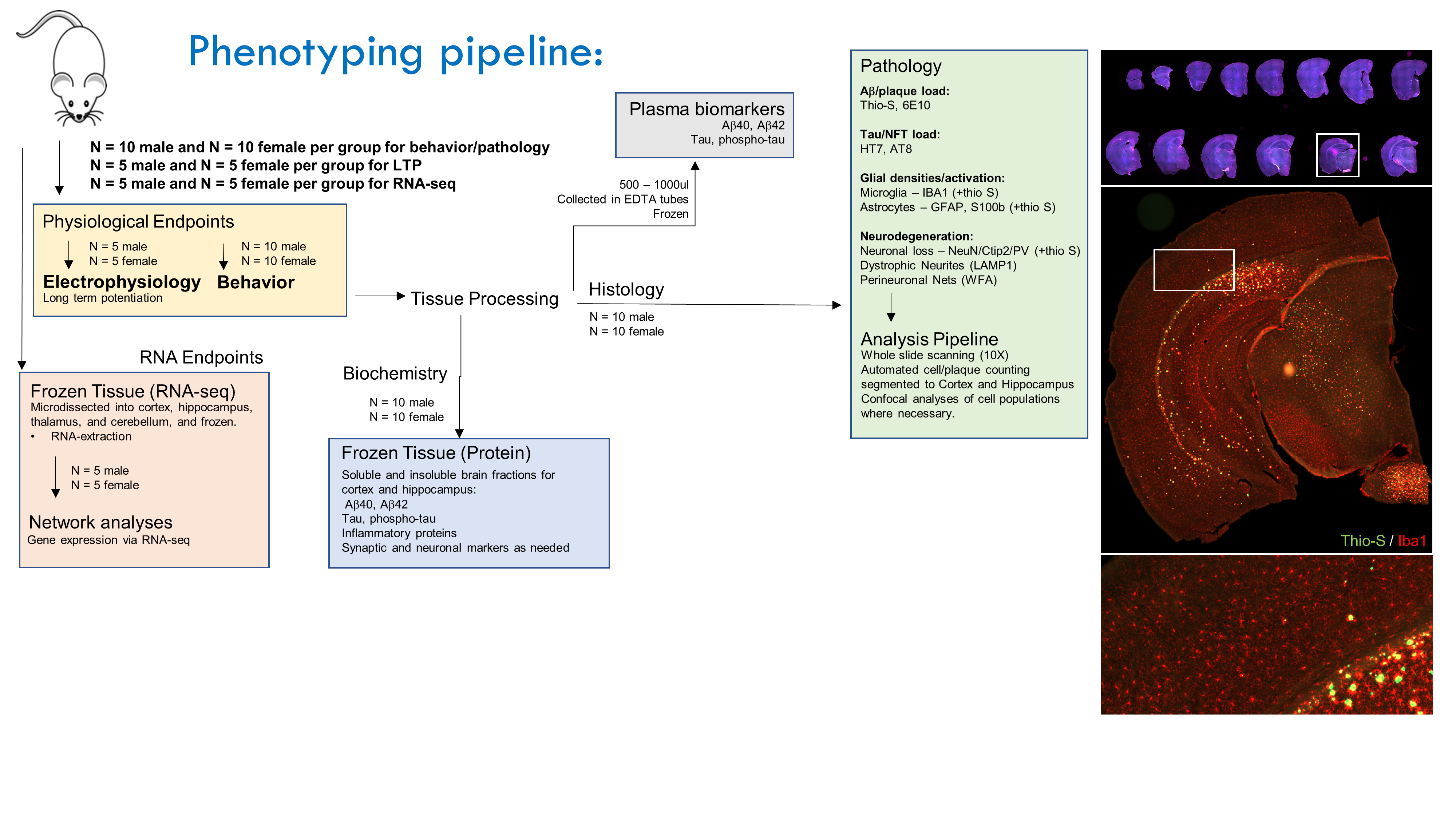
